# The pseudoproteinase iRhom2 critically promotes acute lung inflammation

**DOI:** 10.64898/2026.06.05.730404

**Authors:** Christine Lux, Selcan Kahveci-Türköz, Katharina Schun, Aaron Babendreyer, Christian Martin, Petr Kasparek, Radislav Sedlacek, Stefan Düsterhöft, Andreas Ludwig

## Abstract

ADAM17 sheds cell surface molecules such as TNF-α, IL-6Rα and L-selectin. This activity requires either iRhom1 or iRhom2 as adapter molecules. Since iRhom2 is predominantly expressed in leukocytes and upregulated in tissue cells during inflammation, it represents a potential anti-inflammatory target. We therefore investigated the effects of iRhom2 deficiency in mice using in vivo, ex vivo, and in vitro models of acute inflammation.

In an in vivo model of LPS-induced lung inflammation, iRhom2 knockout mice showed reduced neutrophil recruitment into the bronchoalveolar space. Notably, the few recruited neutrophils remained L-selectin positive, whereas most neutrophils in wildtype mice were L-selectin negative, confirming that L-selectin shedding depends on the iRhom2/ADAM17 axis. Furthermore, it suggests that impaired shedding is associated with decreased neutrophil recruitment.

Additionally, ADAM17-dependent release of TNF-α and IL-6Rα into the alveolar space was diminished in the absence of iRhom2, accompanied by reduced expression of inflammatory mediators. In isolated perfused lungs challenged with LPS, iRhom2 deficiency similarly reduced inflammatory mediator production, indicating a role for iRhom2 in resident lung tissue cells during the initiation of inflammation.

To specifically assess immune cell responses, we further examined macrophages, the sole resident immune cells in the lung. In vitro, LPS-stimulated bone marrow derived macrophages lacking iRhom2 showed decreased shedding of TNF-α and IL-6Rα and reduced induction of secondary inflammatory mediators.

Thus, targeting iRhom2 effectively suppresses ADAM17-mediated inflammatory responses in the lung, while preserving basal ADAM17 activity through iRhom1, offering a more selective therapeutic strategy with fewer side effects.

**Highlights:**

1. The ADAM17 adapter molecule iRhom2 is required for the acute lung inflammation of mice in vivo including cytokine response and neutrophil recruitment.
2. In resident lung tissue cells iRhom2 promotes the LPS induced inflammatory response at the alveolar interface.
3. In macrophages iRhom2 is required for an effective ADAM17 dependent inflammatory response to LPS.
4. Thus, iRhom2 targeting can serve to suppress inflammatory activities of ADAM17 in the lung.

**Graphical Abstract:** Schematic overview depicting the role of the iRhom2-ADAM17 axis in mediating neutrophil infiltration and cytokine release during induced pulmonary inflammation, serving as a model for acute lung injury (ALI).

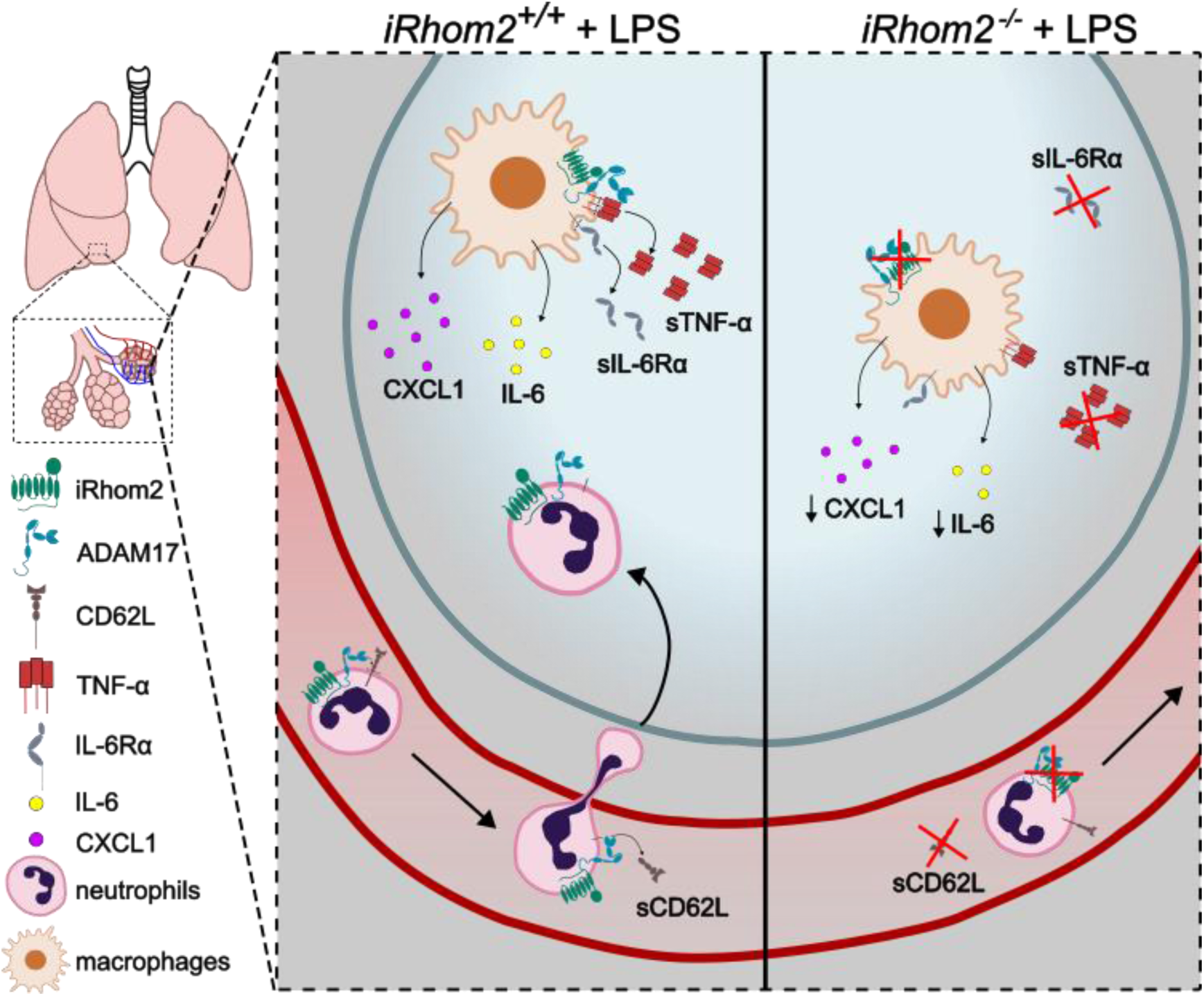

## Introduction

Acute lung injury (ALI) and its severe form, acute respiratory distress syndrome (ARDS), are associated with high morbidity and mortality rates and the only effective therapy so far being supportive mechanical ventilation [1]. ALI and ARDS are characterized by dysregulated acute inflammation, leading to disruption of the alveolar epithelial-endothelial barrier, neutrophil infiltration and the excessive production of proinflammatory mediators (reviewed in [2]). ALI can be triggered by a variety of direct and indirect insults, including pneumonia, aspiration, sepsis and bacterial endotoxins such as lipopolysaccharide (LPS), which activate innate immune responses and initiate inflammatory signaling cascades (reviewed in [2, 3]). Thus, early in the course of ALI, alveolar macrophages orchestrate the inflammatory response through the release of cytokines and chemokines, promoting neutrophil activation and recruitment, which are central to disease progression [4]. Especially, elevated levels of circulating tumor necrosis factor (TNF)-α, interleukin (IL)-6 and IL-8 correlate with disease severity [5].

Circulating TNF-α is believed to act as an initiating cytokine in lung injury development [6]. TNF-α is expressed as a precursor molecule by various cell types but mainly by macrophages. This molecule is converted into a soluble TNF-α by limited proteolysis at the cell surface predominantly mediated by ‘A Disintegrin And Metalloproteinase’ (ADAM)17 [7]. Soluble TNF-α together with other ADAM17 substrates such as IL-6 receptor α (IL-6Rα) and L-selectin (CD62L) were shown to contribute to the LPS-induced septic shock [8–10]. Hence, ADAM17 might play a key role in development of ALI.

Whereas a recent study by Aljohmani et al. suggested that ADAM17 might dampen the local proinflammatory response in the lung [11], the metalloprotease has previously been considered an enhancer of lung injury (reviewed in [12]). Thus, the determination of pro-versus anti-inflammatory responses might not be governed by ADAM17 itself, but rather by its regulation through the pseudoproteinases iRhom1 and iRhom2. While iRhom1 was shown to be generally expressed at higher levels than iRhom2 in non-immune cells, the latter is upregulated upon inflammatory stimulation [13]. Furthermore, iRhom2 is predominantly found in immune cells, where iRhom1 is mainly absent [14, 15]. Accordingly, iRhom2 knockout (*iRhom2^-/-^*) mice display normal phenotypes, in contrast to ADAM17 and iRhom1 knockouts which exhibit lethal phenotypes [15–17]. However, *iRhom2^-/-^*mice lack TNF-α release, indicating impaired ADAM17 maturation in macrophages and other immune cells that depend on iRhom2-mediated ADAM17 surface expression [16]. Consequently, these mice survive doses of LPS that are typically lethal due to TNF-α-mediated sepsis in wildtype mice [18].

In the present study, we investigated the regulation and role of iRhom2 in the early onset of lung inflammation. Database analysis indicated upregulation of iRhom2 under inflammatory conditions in human patients and this was confirmed for LPS-induced inflammation in mice lungs, in isolated perfused lungs and in isolated bone marrow derived macrophages. The functional relevance of iRhom2 was studied in mice lacking iRhom2. Here, iRhom2 deficiency reduced neutrophil infiltration into mice lungs and reduced shedding of cardinal inflammatory ADAM17-substrates including L-selectin, TNF-α and IL-6Rα. Analysis of in vivo, ex vivo and in vitro experiments however indicated that not only these substrates but also secondary inflammatory mediators were reduced in the absence of iRhom2. Our findings therefore highlight a proinflammatory role of iRhom2 in the inflamed lung.

## Materials and Methods

### Bioinformatics

Publicly available single-cell RNA-sequencing data were obtained from GEO dataset GSE196638 [19], comprising three control human lung samples (NL_1, NL_2, NL_3) and three emphysema lung samples (TXP24, TXP25, TXP26). Raw 10x Genomics count matrices were processed in Python 3.11.4 using Scanpy 1.11.5 and AnnData 0.9.2. Samples were imported individually, annotated by donor and condition, and concatenated into a single AnnData object. Raw counts were stored prior to normalization.

Per-cell quality control metrics included the number of detected genes, total counts, and the fractions of mitochondrial, ribosomal, and hemoglobin-associated transcripts. Cells were retained if they contained 300–6000 detected genes, 500–50,000 total counts, and ≤20% mitochondrial counts. Ribosomal and hemoglobin transcript fractions were inspected but not used for global filtering.

After quality control, counts were normalized to 10,000 counts per cell and log1p-transformed. The complete log-normalized expression matrix was retained for downstream visualization and quantification of all detected genes. Highly variable genes (HVGs) were identified using Scanpy’s dispersion-based Seurat-style implementation. Importantly, HVGs were used only for scaling, principal component analysis, neighborhood graph construction, UMAP visualization, and Leiden clustering, whereas all gene-level analyses were performed using the complete log-normalized expression matrix. The top 3000 HVGs were used for dimensionality reduction. A k-nearest-neighbor graph was constructed using the first 30 principal components and 15 nearest neighbors, followed by UMAP embedding and Leiden clustering at a resolution of 0.5.

Clusters were manually annotated based on canonical lung cell type markers and cluster-specific marker genes identified with Scanpy’s Wilcoxon rank-sum test. Cell type proportions were calculated per donor and condition. Genes of interest were quantified across cell types, compartments, conditions, and donors using mean log-normalized expression and the fraction of expressing cells.

### Mice

Animal model experiments were approved by local government authorities and were performed according to the German animal protection law (approval-ID: 81-02.04.2019.A271; A4-90029A4). Mice with iRhom2-knockout (*iRhom2^-/-^*) and wildtype (*iRhom2^+/+^*) littermates in a C57BL/6 background derived from in house breeding [20]. Experiments were performed on 8-12 weeks old mice.

### LPS model of acute lung inflammation

Mice were intranasally instilled with 20 µl per nostril LPS (400 µg/ml body weight in PBS) or PBS as vehicle control. After 4 h or 24 h, bronchoalveolar lavage (BAL) of the left lung was collected with intratracheal administration of 0.5 ml PBS. BAL was centrifuged (400×g, 10 min, 4 °C) and supernatant was used for ELISA measurements of cytokine release while sediment cells were further analyzed by flow cytometry. The right superior lobe was filled with 4 % paraformaldehyde (PFA) for 5 min and further processed for histological analysis as described previously [21]. The middle and inferior right lobes were used for RNA and protein analysis.

### Isolated perfused lung (IPL)

For IPLs, mice were prepared as previously described [22]. They were ventilated with a respiratory rate of 90 breaths per minute by a negative pressure from -3 cm to -8 cm H_2_O in order to have a tidal volume of ∼200-250 µl. Every 5 minutes, a hyperinflation of -20 cm H_2_O was performed. A previously described Gelafundin^®^ ISO 40 mg/ml (Braun) supplemented medium was used for perfusion [23]. After 30 min of baseline measurements, 50 µl LPS (400 µg/kg body weight in PBS) or PBS (vehicle control) were administered intratracheally with a microsprayer (PennCentury). Afterwards, mice were ventilated as described for another 180 min. Data analysis was performed with the Pulmodyn software (Hugo Sachs). The mice lungs were then detached from the mouse body and the accessory lobe (lobus accessorius pulmonis) was separated and used to analyze edema formation with the wet-to-dry (W/D) ratio. The bronchoalveolar lavage (BAL) was collected after intratracheal administration of 1 ml PBS. Lung tissue was further processed for RNA and protein extraction.

### Wet-to-dry (W/D) ratio

To determine edema formation, the wet weight of the accessory lobe of the right lung was immediately determined after finishing the IPL. The tissue was then dried at 60 °C for 72 h and weighed again to obtain its dry weight. The W/D ratio was calculated by dividing the wet weight through the dry weight.

### Bone marrow derived macrophages (BMDMs)

BMDMs were isolated from femur and tibia of mice hind legs as described before [20]. At day 10, differentiated BMDMs were seeded on 12-well plates. On the next day, cells were stimulated with 1 ng/ml LPS or vehicle control for 0.5, 1, 2, 4, 8 or 24 h. Supernatants were used for ELISA and cells were lysed for qPCR analysis.

### Flow cytometry (FC)

BAL cells from the in vivo LPS-induced acute lung inflammation model were fixed with 1 % PFA in FC buffer (PBS with 0.1% BSA and 1mM EDTA) for 5 min in the dark. Cells were then centrifuged (400×g, 10 min, 4 °C) and resuspended in FC buffer. Cells were further processed as described before [24]. In this study, following antibodies were used for differentiation of immune cell types: FITC rat anti-mF4/80, clone Cl:A3-1 (MCA497F, BioRad); APC rat anti-mLy6G, clone 1A8 (130-123-564, Miltenyi Biotec); PE rat anti-mCD62L, clone MEL-14 (12-0621-82, Invitrogen); eFluor^TM^ 450 rat anti-mCD11b, clone M1/70 (48-0112-82, Invitrogen); APC-Cy^TM^7 hamster anti-mCD11c, clone HL3 (561241, BD Pharmingen); V500 syrian hamster anti-mCD3e, clone 500A2 (560773, BD Horizon); VioBlue^®^ rat anti-mCD45R, clone RA3-6B2 (120-004-170, Miltenyi Biotec).

### Enzyme-linked immunosorbent assay (ELISA)

Supernatants from BMDMs and BAL from acute lung inflammation model or IPL experiments were centrifuged at 16’000×g for 5 min at 4 °C to remove cell debris. Commercial DuoSet ELISA Kits from R&D Systems were used according to manufacturer’s instructions to detect soluble TNF-α (cat.#:DY410), IL-6 (cat.#:DY406), IL-6Rα (cat.#:DY1830) and CXCL1/KC (cat.#:DY453) from the supernatants and BAL. The substrate BM Blue POD (Roche) was used to start the reaction which was stopped with 5 % H_2_SO_4_. Background signal measured at 550 nm with SpectraMax iD3 microplate reader (Molecular Devices, LLC) was subtracted from the absorbance measured at 450 nm to quantify protein concentrations.

### Western blot

For protein analysis, ∼30 mg lung tissue was processed with the Mixer Mill MM 400 (Retsch) in 300 µl protein lysis buffer (20mM Tris, 150 mM NaCl, 4 mM EDTA, 1 mM PMSF, 10 µM GI254023X, 1 % Triton X-100) supplemented with cOmplete^TM^ protease inhibitor. Cells from BMDMs were lysed with 150 µl protein lysis buffer (50 mM Tris, 134 mM NaCl, 2 mM EDTA, 1% Triton X-100, 1x cOmplete^TM^ protease inhibitor (Merck), 10 mM 1,10-phenanthroline, 1 mM Na_3_VO_4_, 10 mM β-Glycerophosphate, 30 mM NaF, pH=7.5). After centrifugation of the samples for 10 min and 16,000×g at 4 °C, laemmli buffer (520 mM SDS, 40 % (v/v) 250 mM Tris (pH7.5), 35 % (v/v) glycerol, 25 % (v/v) β-mercaptoethanol, 3 mM bromphenol blue) was added to the supernatants for protein denaturation. Protein samples were separated in an SDS-PAGE and transferred to a polyvinylidene difluoride (PVDF) membrane with a pore size of 0.45 µm (Millipore, Immobilon-FL). Membranes were blocked for 30 min with 3 % (w/v) BSA in 0.1 % Tween-TBS (50 mM Tris, 150 mM NaCl, pH 7.4) and incubated overnight at 4 °C with primary antibodies. In this study, the used primary antibodies were diluted as follows: rabbit anti-ADAM17 (1:1666 in 1 % w/v BSA in 0.1 % Tween-TBS, ab39162, Abcam), rabbit anti-iRhom2 (1:1000 in 1 % w/v BSA and 3 % w/v non-fat dry milk in 0.1 % Tween-TBS, SAB1304414, Sigma-Aldrich), mouse anti-GAPDH (1:2000 in 1 % w/v BSA in 0.1 % Tween-TBS, MA5-15738, ThermoFisher), mouse anti-STAT3 (1:1000 in 1 % w/v BSA in 0.1 % Tween-TBS, 9139, Cell Signaling), rabbit anti-phospho-STAT3 (1:1000 in 1 % w/v BSA in 0.1 % Tween-TBS, 9131, Cell Signaling), rabbit anti-NFκB p65 (1:1000 in 1 % w/v BSA in 0.1 % Tween-TBS, 8242, Cell Signaling), rabbit anti-phospho-NFκB p65 (1:1000 in 1 % w/v BSA in 0.1 % Tween-TBS, 3033, Cell Signaling). The secondary antibodies DyLight^TM^ 680-conjugated goat anti-mouse (ThermoFisher; 35519) and Alexa Fluor^TM^ Plus 800-conjugated anti-rabbit (ThermoFisher; A32808) diluted 1:100,000 in 1 % w/v BSA in 0.1 % Tween-TBS were added for 1.5 h at room temperature. Before fluorescence detection with the Odyssey 9120 157 imager system (LI-COR), the membrane was washed once with 0.1 % Tween-TBS and twice with TBS. Band intensities were quantified using Image Studio Lite software v5.2 (LI-COR). To normalize the biological replicates, the band intensity of each sample was divided by the total of sample values from the same membrane (sum of all data points in a replicate) [25].

### Quantitative real-time polymerase chain reaction (qPCR)

Previous to RNA extraction, 15-20 mg lung tissue were processed in 600 µl buffer RLT (Qiagen) containing 1 % β-mercaptoethanol with the Mixer Mill MM 400 (Retsch) or 500.000 cells were lysed with 350 µl buffer RLT (Qiagen). The RNA extraction was then performed using the RNeasy Kit (Qiagen) followed by a photometric concentration measurement with Nanodrop (Peqlab). 1000 ng RNA of lung tissue from the acute lung inflammation model, 750 ng RNA of lung tissue from isolated perfused lung model and 300 ng RNA from bone marrow derived macrophages were reversed transcribed using the PrimeScript^TM^ RT Reagent Kit (Takara Bio Europe) following manufacturer’s instructions. The qPCR was performed using the iTaq^TM^ Universal SYBR^®^ Green Supermix (Bio-Rad) in the CFX Connect Real-Time PCR Detection System (Bio-Rad). The used forward and reverse primers with the appropriate annealing temperatures are listed in Table 1. Following settings were defined in the used protocol: initial denaturation (95 °C, 5 min), 39 cycles of denaturation (95 °C, 10 s), annealing (temperature in Table 1, 30 s) and amplification (72 °C. 15 s).

**Table 1:**
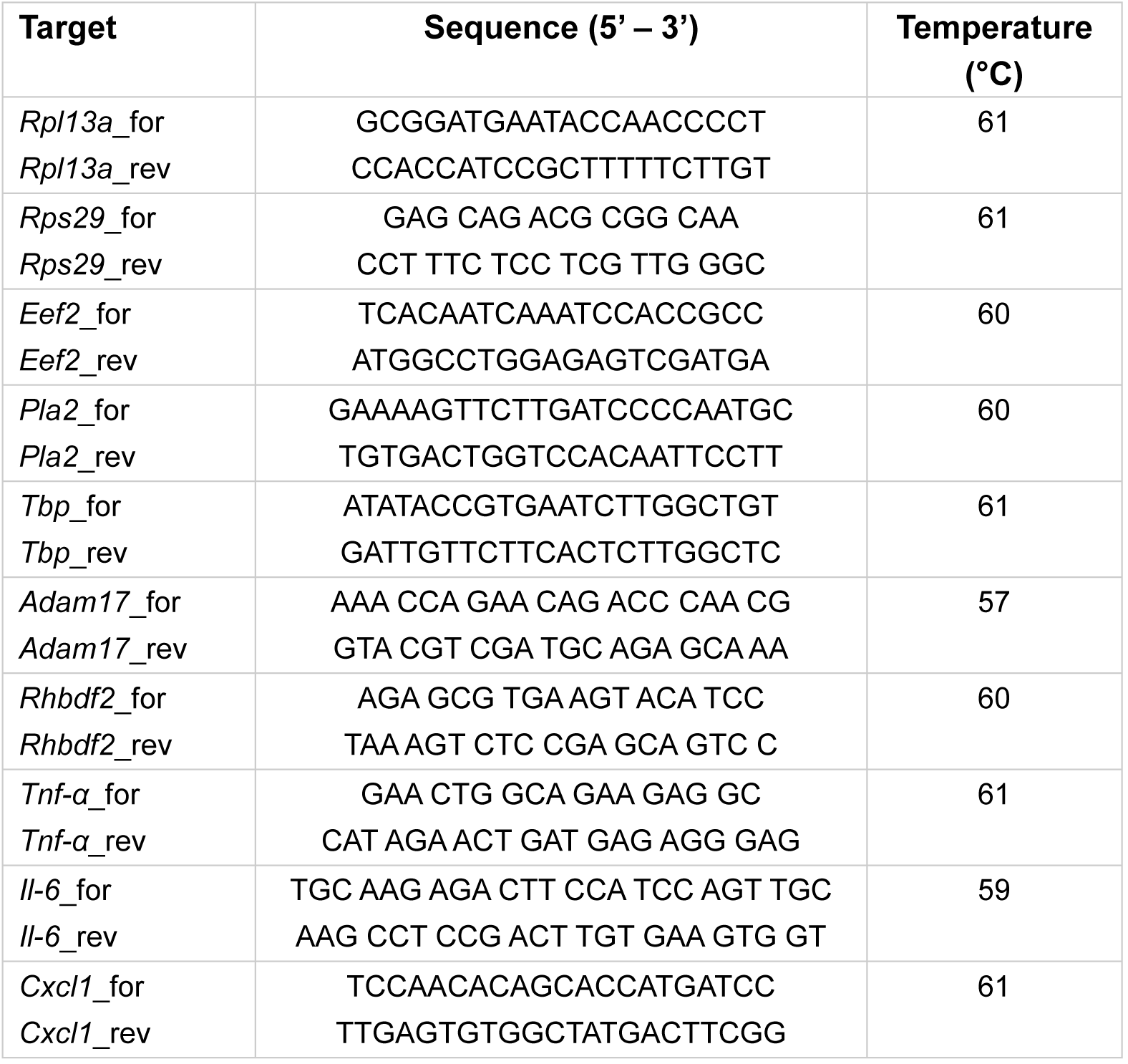
Primers used for qPCR with corresponding sequences and annealing temperatures.

LinRegPCR version 2020.0 software was used to determine the qPCR efficiency with uncorrected RFU values. The quantification was performed with the CFX Maestro Software 2.3 (Bio-Rad) relative to the reference genes *Rpl13a* and *Rps29* for IPL tissue and BMDMs or *Eef2*, *Pla2* and *Tbp* for acute lung inflammation model tissue. The reference genes were defined after reference gene analysis via CFX Maestro Software 2.3 (Bio-Rad).

### Statistics

All experiments were independently repeated at least three times, as indicated in the figure legends. Quantitative data are presented as mean ± standard deviation (SD). Statistical evaluation was performed using a generalized mixed model (PROC GLIMMIX, SAS 9.4; SAS Institute Inc.), with the day of the experiment included as a random effect to account for inter-experimental variability. The distribution of the data (normal, lognormal) was assessed, and diagnostic tests-including residual analysis and the Shapiro-Wilk test were applied to evaluate model assumptions. In cases of heteroscedasticity, as identified using the COVTEST statement, the degrees of freedom were adjusted using the Kenward-Roger approximation. To control for multiple testing, p-values were adjusted using the false discovery rate (FDR) method. Outliers were excluded with ROUT method (Q=1%). A p-value of <0.05 was considered statistically significant. Statistical significance is denoted as follows: * p<0.05, ** p<0.01, *** p<0.001. All figures were generated using GraphPad Prism version 8 (GraphPad Software).

## Results

### iRhom2 is upregulated in macrophages under inflammatory conditions

To gain insight into the role of iRhom2 in lung tissue, single cell mRNA sequencing data from healthy human lungs were reanalyzed. As anticipated, *RHBDF2* which encodes for iRhom2 was predominantly expressed in alveolar and inflammatory macrophages as well as in inflammatory neutrophil-like cells, while *RHBDF1* encoding for iRhom1 was more abundant in other lung tissue cell populations (Figure 1 a-c). In the same study, lung tissue biopsies from patients with emphysema showed increased macrophage *RHBDF2* mRNA expression compared to healthy controls (Suppl. Figure S 1) suggesting a potential role in lung-damaging inflammatory processes.

**Figure 1:**
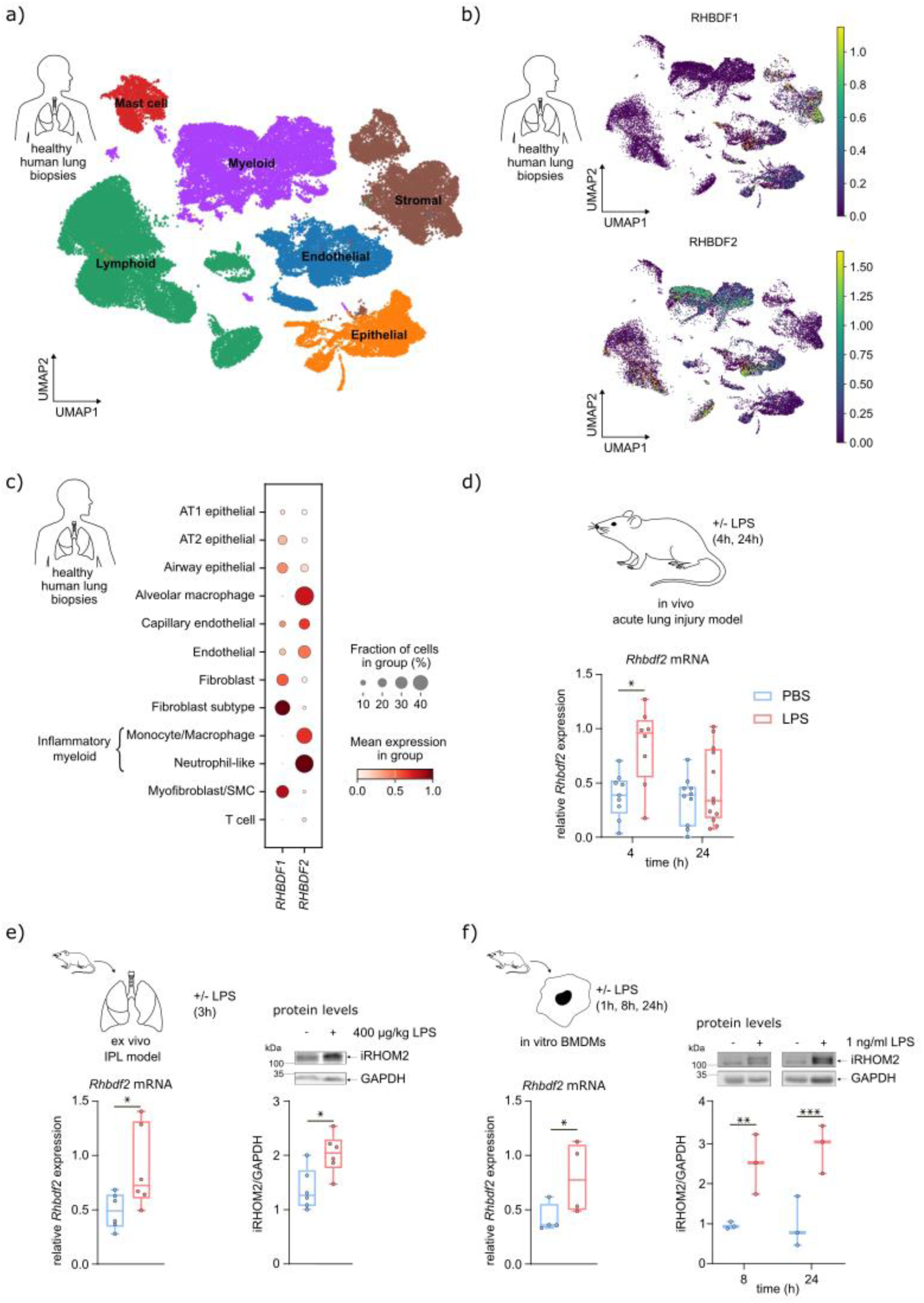
iRhom2 is upregulated under inflammatory conditions. **a)** UMAP plot visualization of lung from healthy lung biopsies in 6 clusters. Data were reanalyzed from GSE196638. **b)** UMAP plot showing *RHBDF1* (encoding for iRhom1) and *RHBDF2* (encoding for iRhom2) are differentially expressed in different lung cell types. **c)** Single cell mRNA expression of *RHBDF1* and *RHBDF2* in healthy lung biopsies. AT1/2 – alveolar type 1/2; SMC – smooth muscle cell. **d)** Quantified relative *Rhbdf2* mRNA expression measured by qPCR analysis of processed wildtype mice lung tissue 4 h and 24 h following 400 µg/kg body weight LPS or identical volume of PBS administration. **e)** Quantified relative *Rhbdf2* mRNA expression measured by qPCR analysis and Western blot analysis of iRHOM2 protein in processed wildtype mice lung tissue deriving from isolated perfused lung experiments following 400 µg/kg body weight LPS or identical volume of PBS administration. GAPDH serves as a loading control for Western blot analysis. **f)** Quantified relative *Rhbdf2* mRNA expression measured by qPCR analysis 1 h following 1 ng/ml LPS addition and Western blot analysis of iRHOM2 protein from bone marrow derived macrophages following 8 h and 24 h following 1 ng/ml LPS treatment. GAPDH serves as a loading control for Western blot analysis. **d-f)** Quantitative data are shown as Tukey box plots from at least four independent experiments. Statistical analysis was performed using a generalized linear mixed model with a post-hoc false discovery rate (FDR) correction for multiple comparisons. Statistical differences are indicated by bars and asterisks with * p ≤ 0.05, ** p ≤ 0.01 and *** p ≤ 0.001, n.s. *–* not significant.

To further investigate this, the role of iRhom2 in ALI was assessed. Since no suitable mRNA sequencing data were available from ALI or ARDS patients, lung tissue from mice following LPS inhalation was analyzed by qPCR. Murine *Rhbdf2* mRNA was already increased 4 h after induction of inflammation (Figure 1 d). Moreover, inflammation-associated upregulation of iRhom2 was observed on mRNA and protein levels in the isolated perfused lung (IPL) model, which is devoid of circulating neutrophils (Figure 1 e). Together with single cell mRNA sequencing data, these findings suggest that macrophages are likely a major source of increased iRhom2 expression. Consistently, LPS stimulation of isolated bone marrow derived macrophages (BMDMs) resulted in a significant upregulation of iRhom2 at both mRNA and protein levels (Figure 1 f).

Collectively, these data suggest that macrophage-derived iRhom2 may contribute to lung inflammatory processes.

### iRhom2 is required for effective neutrophil recruitment during acute lung inflammation

To assess the functional relevance of iRhom2, previously described mice lacking iRhom2 protein were studied in the model of LPS induced lung inflammation (Figure 2 a). Since iRhom2 protein levels could not be assessed due to lack of appropriate antibody for full lung tissue, we confirmed iRhom2 deficiency at the level of ADAM17 maturation (Figure 2 b). As expected, maturation of ADAM17 (mADAM17) from its inactive precursor pADAM17 was significantly reduced in *iRhom2^-/-^* mice, emphasizing the importance of iRhom2 for ADAM17 function in lung tissue. This iRhom2-mediated effect on ADAM17 maturation was observed in untreated as well as LPS-stimulated mice.

**Figure 2:**
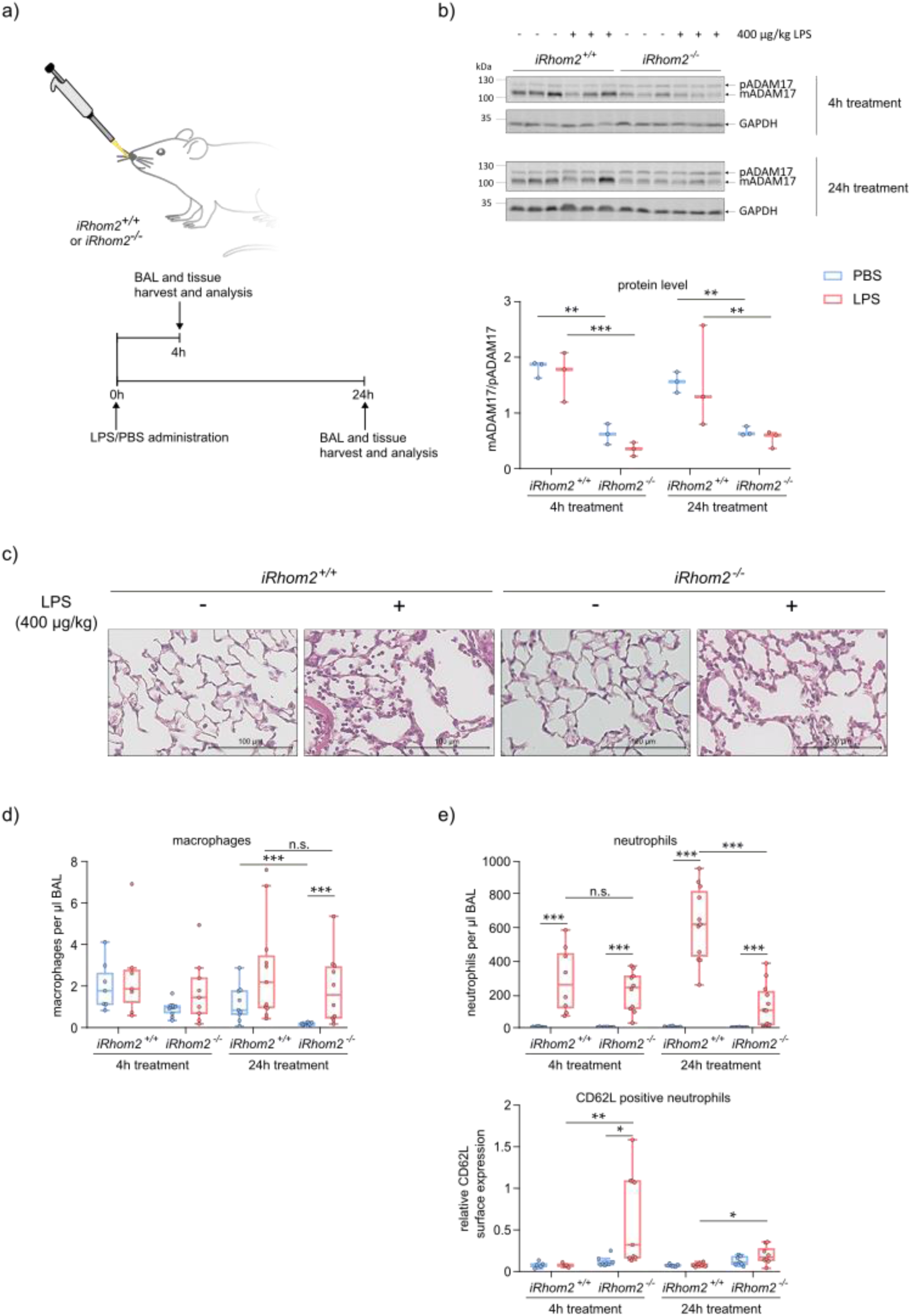
iRhom2 deficiency reduces pulmonary neutrophil infiltration in acute lung inflammation. **a)** Schematic overview of the in vivo acute lung inflammation model. **b)** Western blot analysis of mature ADAM17 (mADAM17) and its proform pADAM17 in processed *iRhom2^+/+^* and *iRhom2^-/-^* lung tissue 4 h and 24 h following 400 µg/kg body weight LPS or identical volume of PBS administration. GAPDH serves as a loading control. Quantification of the Western blots was performed with the Image Studio Lite software v5.2 (LI-COR) and the ‘sum of all data points in a replicate’ method [25]. Quantified data show the determined protein ratio of mADAM17 and the corresponding pADAM17. **c)** 24 h after 400 µg/kg body weight LPS or identical volume of PBS administration, 3 µm-sections of formalin-fixed and paraffin-embedded lung tissue were stained with hematoxylin-eosin. Representative images are shown. **d)** Quantification of macrophages per µl BAL from *iRhom2^+/+^* and *iRhom2^-/-^* mice 4 h and 24 h following 400 µg/kg body weight LPS or identical volume of PBS administration analyzed by flow cytometry. **e)** Quantification of neutrophils per µl BAL from *iRhom2^+/+^* and *iRhom2^-/-^* mice as well as their CD62L surface expression 4 h and 24 h after 400 µg/kg body weight LPS or identical volume of PBS administration analyzed by flow cytometry. **b-e)** Quantitative data are shown as Tukey box plots from at least four independent experiments. Statistical analysis was performed using a generalized linear mixed model with a post-hoc false discovery rate (FDR) correction for multiple comparisons. Statistical differences are indicated by bars and asterisks with * p ≤ 0.05, ** p ≤ 0.01 and *** p ≤ 0.001, n.s. – not significant.

Histological analysis of lung sections revealed that LPS treatment led to increased cellular infiltration in the lung tissue and bronchoalveolar space of wildtype mice, whereas this appeared to be reduced in *iRhom2^-/-^*mice (Figure 2 c). As macrophages represent the primary immune cell population in healthy lung tissue, their number was determined in the bronchoalveolar space by lung lavage and subsequent flow cytometry analysis. While basal levels of macrophages differed between wildtype and knockout mice 24 h following PBS administration, macrophage recruitment was comparable between *iRhom2^-/-^* and wildtype mice 24 h after LPS administration (Figure 2 d). In contrast, neutrophil levels increased significantly after just 4 h following LPS administration compared to PBS control. At 24 h, neutrophil recruitment was then significantly reduced in *iRhom2^-/-^* mice compared to their wildtype littermates (Figure 2 e). Furthermore, the few neutrophils recruited in *iRhom2^-/-^* mice were predominantly L-selectin (CD62L)-positive while the majority of neutrophils in wildtype mice did not express CD62L (Figure 2 e).

These findings confirm a role of the iRhom2/ADAM17 axis in regulating CD62L shedding and indicate the importance of a functional iRhom2/ADAM17 complex formation for neutrophil recruitment to the lung.

### iRhom2 deficiency reduces the cytokine response during acute lung inflammation

Besides CD62L, also TNF-α and IL-6Rα undergo proteolytic shedding. Therefore, the expression and release of both TNF-α and IL-6Rα were studied at mRNA and protein level. While *Tnf-α* mRNA expression increased significantly upon LPS stimulation in both genotypes, TNF-α protein release was undetectable in *iRhom2^-/-^* mice at all time points (Figure 3 a) which is in agreement with ADAM17 being the predominant TNF-α sheddase [26]. Additionally, TNF-α release in wildtype mice was more prominent at 4 h than at 24 h, confirming TNF-α’s role in initiating acute lung inflammation. In contrast, genotype differences in the release of IL-6Rα became apparent only at 24 h after LPS administration (Figure 3 b). In addition to the direct substrates of ADAM17-mediated shedding, we also studied regulation of secreted inflammatory mediators including IL-6 and CXCL1 (also known as KC). The expression of *Il-6* and *Cxcl1* was upregulated upon LPS administration in both genotypes (Figure 3 c-d). Notably, CXCL1 release was significantly reduced in the BAL of *iRhom2^-/-^* mice 4 h after lung inflammation induction, whereas IL-6 release was reduced at 24 h.

**Figure 3:**
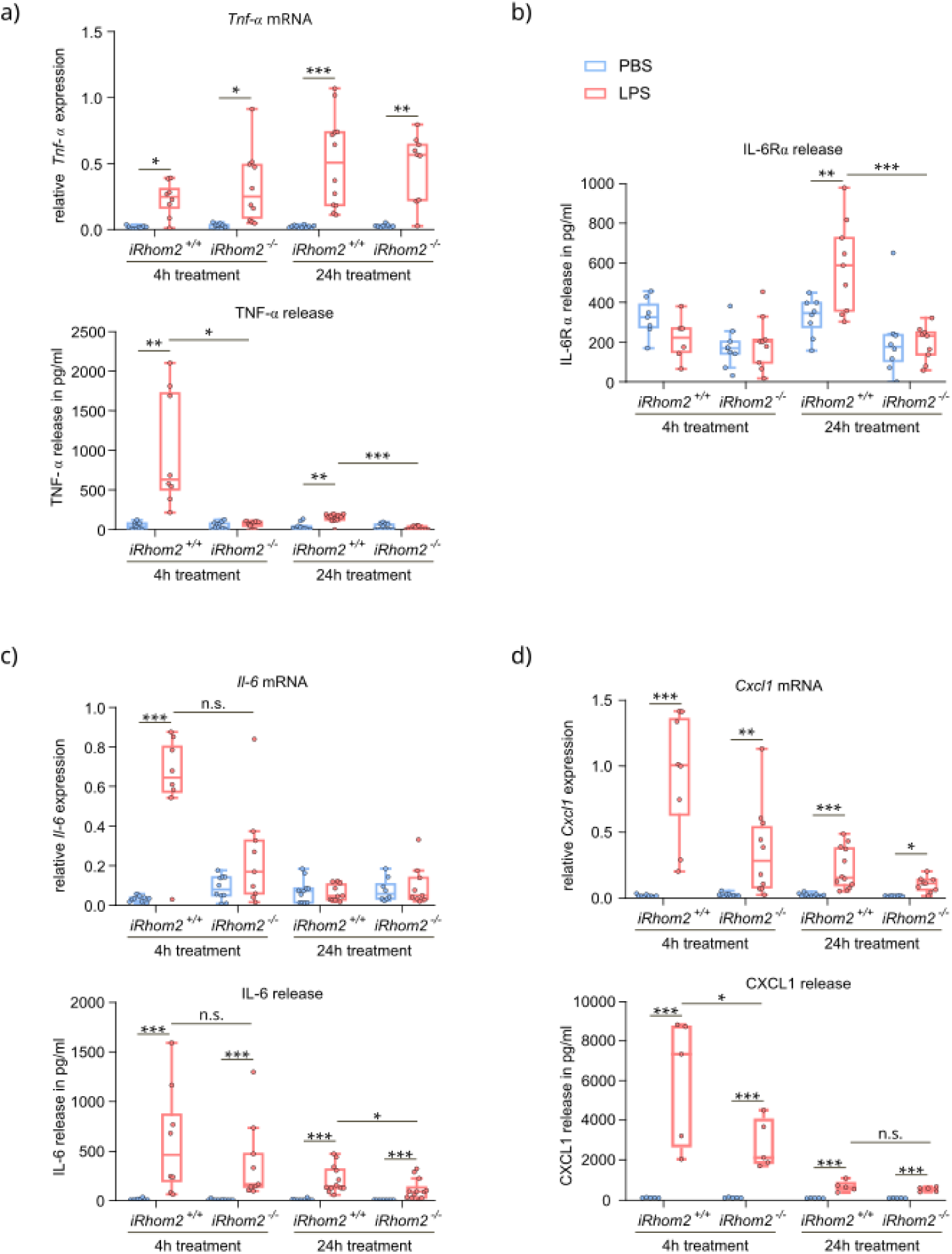
iRhom2 deficiency reduces cytokine and chemokine release in inflamed lungs. **a)** Quantified relative *Tnf-α* mRNA expression measured by qPCR analysis of processed lung tissue and TNF-α protein release in pg/ml determined by ELISA measurements of BAL. **b)** Quantified IL-6Rα release in pg/ml determined by ELISA measurements of BAL. **c)** Quantified relative *Il-6* mRNA expression measured by qPCR analysis of processed lung tissue and IL-6 protein release in pg/ml determined by ELISA measurements of BAL. **d)** Quantified relative *Cxcl1* mRNA expression measured by qPCR analysis of processed lung tissue and CXCL1 protein release in pg/ml determined by ELISA measurements of BAL. **a-d)** Quantitative data are shown as Tukey box plots from at least five independent experiments. Statistical analysis was performed using a generalized linear mixed model with a post-hoc false discovery rate (FDR) correction for multiple comparisons. Statistical differences are indicated by bars and asterisks with * p ≤ 0.05, ** p ≤ 0.01 and *** p ≤ 0.001, n.s. – not significant.

Together, these data indicate that iRhom2 directly contributes to the release of the substrates TNF-α and IL-6Rα, but also indirectly promotes expression and release of the key soluble mediators IL-6 and CXCL1 during acute inflammation.

### iRhom2 promotes the LPS-induced inflammatory response of resident lung tissue cells at the alveolar interface

To investigate the local inflammatory response of resident lung tissue cells independently of recruited neutrophils, an isolated perfused lung model was employed (Figure 4 a). Although iRhom2 deficiency led to significantly decreased ADAM17 maturation, the upregulation of iRhom2 in LPS-treated wildtype mice was associated with a reduced mADAM17/pADAM17 protein ratio (Figure 4 b-c), which may reflect increased turnover of the mature form after its activation [27]. To assess edema formation, the wet-to-dry ratio of the lung was determined. Wildtype mice exhibited increased edema formation following LPS administration compared to *iRhom2^-/-^* mice, indicating enhanced tissue permeability (Figure 4 d). PBS-treated controls showed comparable wet-to-dry ratio in wildtype and knockout mice (Suppl. Figure S 2 a). As observed in the in vivo model, mRNA expression of *Tnf-α, Il-6* and *Cxcl1* increased to a similar extent in both genotypes following LPS administration (Figure 4 e,g,h). As expected, TNF-α protein release was detected only in LPS-treated wildtype mice but not in *iRhom2^-/-^*mice (Figure 4 e). IL-6Rα release was significantly lower in LPS-treated *iRhom2^-/-^*mice compared to wildtype mice (Figure 4 f).

**Figure 4:**
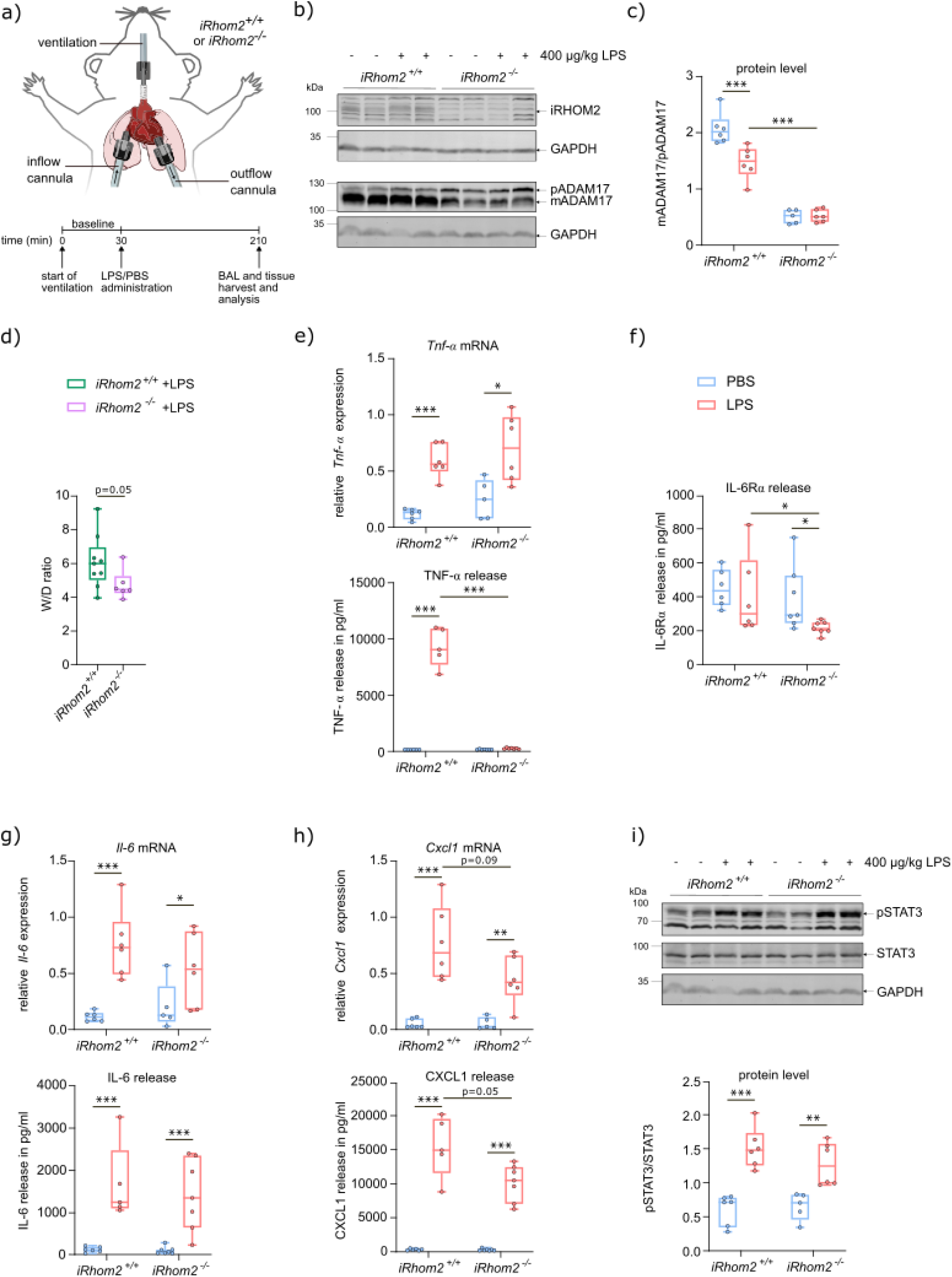
iRhom2 deficiency attenuates proinflammatory mediator release in the isolated perfused lung model. **a)** Schematic overview of the isolated perfused lung (IPL) model. **b)** Western blot analysis of iRhom2, mADAM17 and pADAM17 in processed *iRhom2^+/+^* and *iRhom2^-/-^* lung tissue following 400 µg/kg body weight LPS or identical volume of PBS administration. GAPDH serves as a loading control. **c)** Quantified Western blot data show the determined protein ratio of mADAM17 and the corresponding pADAM17. **d)** Wet-to-dry (W/D) ratio of IPL-derived lungs of *iRhom2^+/+^* and *iRhom2^-/-^* mice subjected to 400 µg/kg body weight LPS. The ratio was calculated by dividing the lung mass dried for 72 h at 60 °C by their wet mass immediately after the experiment. **e)** Quantified relative *Tnf-α* mRNA expression measured by qPCR analysis of processed lung tissue and TNF-α protein release in pg/ml determined by ELISA measurements of BAL. **f)** Quantified IL-6Rα release in pg/ml determined by ELISA measurements of BAL. **g)** Quantified relative *Il-6* mRNA expression measured by qPCR analysis of processed lung tissue and IL-6 protein release in pg/ml determined by ELISA measurements of BAL. **h)** Quantified relative *Cxcl1* mRNA expression measured by qPCR analysis of processed lung tissue and CXCL1 protein release in pg/ml determined by ELISA measurements of BAL. **i)** Western blot analysis of phosphorylated STAT3 (pSTAT3) and STAT3 from processed lung tissue. GAPDH serves as a loading control. Quantified data show the determined protein ratio of pSTAT3 and the corresponding STAT3. **c-i)** Quantitative data are shown as Tukey box plots from at least three independent experiments. Statistical analysis was performed using a generalized linear mixed model with a post-hoc false discovery rate (FDR) correction for multiple comparisons. Statistical differences are indicated by bars and asterisks with * p ≤ 0.05, ** p ≤ 0.01 and *** p ≤ 0.001, n.s. – not significant.

In parallel, the effect on soluble inflammatory mediators was studied. While the release of CXCL1 appeared to be reduced in *iRhom2*^-/-^ mice, no clear differences in IL-6 release were detectable after 3 h of ventilation (Figure 4 g-h). Due to technical limitations, prolonged ventilation of the mouse was not feasible, thus precluding the detection of potential later differences in IL-6 release. Hence, although STAT3 phosphorylation increased following LPS-administration, genotype-related differences in the activation of IL-6-mediated pathway could not be determined (Figure 4 i). The phosphorylation of NFκB downstream of LPS-mediated signaling could not be detected at this time point (Suppl. Figure S 2 b)

Collectively, these findings suggest that iRhom2 modulates the local inflammatory response of resident lung tissue cells.

### iRhom2 is required in macrophages for an effective ADAM17 dependent inflammatory response to LPS

As macrophages constitute the resident immune cell population in lung tissue and orchestrate the primary immune response, bone marrow derived macrophages (BMDMs) were isolated and differentiated for further analysis (Figure 5 a). As expected, maturation of ADAM17 was detected only in wildtype cells and 8 h of LPS stimulation led to a reduced mADAM17/pADAM17 protein ratio compared to control treatment (Figure 5 b-c). Based on findings by Skurski et al. that low concentrations of LPS trigger an iRhom2-dependent macrophage response through an amplification loop via the ADAM17-mediated release of TNF-α [28], a range of LPS concentrations was evaluated in our experiments (Suppl. Figure S 3 a). Concentrations at or above 1 ng/ml LPS showed an induction in cytokine expression (*Tnf-α, Il-6, Cxcl1*) in wildtype compared to *iRhom2*^-/-^ cells under our experimental conditions. Therefore, an LPS concentration of 1 ng/ml was selected for subsequent experiments to allow detection of potential iRhom2-dependent effects under low-dose conditions. As a direct target of ADAM17, TNF-α was released from wildtype BMDMs after 2 h of LPS stimulation and remained detectable throughout the 24 h observation period, whereas it was absent in iRhom2-deficient cells (Figure 5 d, Suppl. Figure S 3 b). Also an iRhom2-dependent induction of IL-6Rα release could be observed after 24 h of LPS stimulation (Figure 5 e).

**Figure 5:**
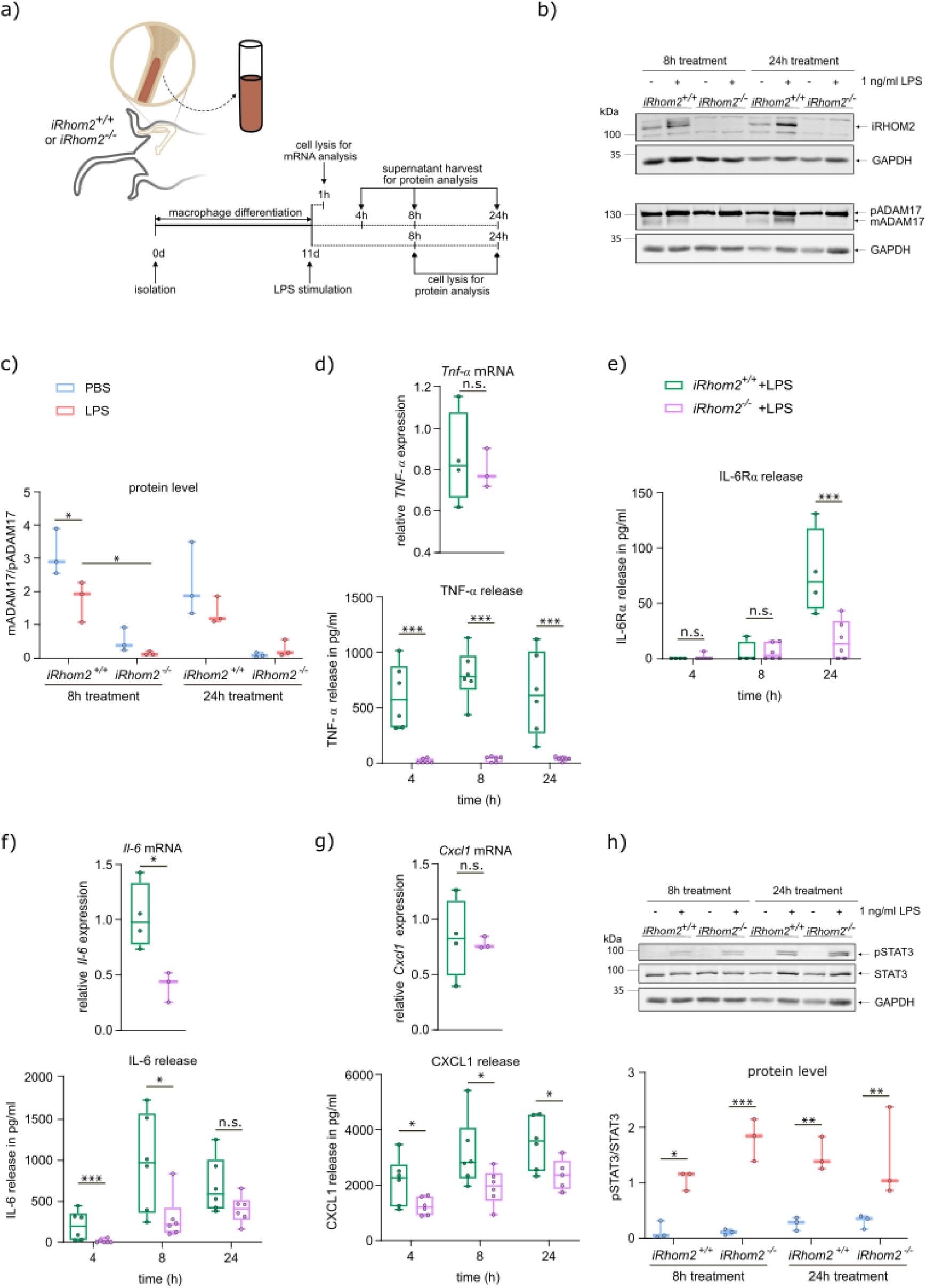
iRhom2 deficiency reduces mediator release from activated macrophages. **a)** Schematic overview of bone marrow derived macrophages (BMDMs) isolation, differentiation and processing. **b)** Western blot analysis of iRhom2, mADAM17 and pADAM17 in *iRhom2^+/+^* and *iRhom2^-/-^*BMDMs 8 h and 24 h following 1 ng/ml LPS or identical volume of PBS stimulation. GAPDH serves as a loading control. **c)** Quantified data show the determined protein ratio mADAM17 and the corresponding pADAM17. **d)** Quantified relative *Tnf-α* mRNA expression measured by qPCR analysis of BMDMs treated for 1 h with 1 ng/ml LPS and TNF-α protein release in pg/ml determined by ELISA measurements of supernatant from BMDMs treated for 4, 8 and 24 h with 1 ng/ml LPS. **e)** Quantified IL-6Rα release in pg/ml determined by ELISA measurements of supernatant from BMDMs treated for 4, 8 and 24 h with 1 ng/ml LPS. **f)** Quantified relative *Il-6* mRNA expression measured by qPCR analysis of BMDMs treated for 1 h with 1 ng/ml LPS and IL-6 protein release in pg/ml determined by ELISA measurements of supernatant from BMDMs treated for 4, 8 and 24 h with 1 ng/ml LPS. **g)** Quantified relative *Cxcl1* mRNA expression measured by qPCR analysis of BMDMs treated for 1 h with 1 ng/ml LPS and Cxcl1 protein release in pg/ml determined by ELISA measurements of supernatant from BMDMs treated for 4, 8 and 24 h with 1 ng/ml LPS. **h)** Western blot analysis of phosphorylated STAT3 (pSTAT3) and STAT3 from BMDMs 8 h and 24 h following 1 ng/ml LPS or identical volume of PBS stimulation. GAPDH serves as a loading control. Quantified data show the determined protein ratio of pSTAT3 and the corresponding STAT3. **c-h)** Quantitative data are shown as Tukey box plots from at least three independent experiments. Statistical analysis was performed using a generalized linear mixed model with a post-hoc false discovery rate (FDR) correction for multiple comparisons. Statistical differences are indicated by bars and asterisks with * p ≤ 0.05, ** p ≤ 0.01 and *** p ≤ 0.001, n.s. – not significant.

We also studied the expression and release of other secreted inflammatory mediators. On the mRNA level, expression of *Il-6* showed iRhom2-dependency similar to that reported by Skurski et al. (Figure 5 f), while *Cxcl1* expression seemed not to be affected (Figure 5 g, Suppl. Figure S 3 a). Differences were more apparent on the protein level, where IL-6 was released in an iRhom2-dependent manner after 4 h and 8 h (Figure 5 f, Suppl. Figure S 3 b) and CXCL1 showed iRhom2-dependent release from 4 h onwards (Figure 5 g, Suppl. Figure S 3 b). Although STAT3 phosphorylation increased at 8 h and 24 h following LPS stimulation, it did not differ between stimulated wildtype and *iRhom2^-/-^*BMDMs, despite iRhom2-dependent release of IL-6 and IL-6Rα (Figure 5 h).

Overall, these data indicate that iRhom2 regulates cytokine and chemokine release in activated macrophages in a time-dependent manner.

## Discussion

Acute lung injury (ALI) and acute respiratory distress syndrome (ARDS) are severe inflammatory conditions associated with high morbidity and mortality. A key hallmark of both conditions is pulmonary neutrophilia, which is considered a major driver of tissue damage (reviewed in [29]). However, the mechanisms governing neutrophil recruitment to the lung remain incompletely understood. Here, we identify iRhom2 as a critical regulator of pulmonary neutrophil infiltration. We showed that loss of iRhom2, and the resulting impairment in ADAM17-mediated L-selectin shedding, markedly reduced neutrophil recruitment to the lung. Consistent with previous findings, this effect was specific to neutrophils, while macrophage infiltration remained unaffected [30]. L-selectin is a cell adhesion molecule abundant on circulating neutrophils and is essential for tethering and rolling during the adhesion cascade that precedes transendothelial migration (TEM) at sites of inflammation. A previous study has shown that L-selectin shedding following its clustering with the endothelial adhesion molecule PECAM-1 enhances the speed of TEM [31]. However, the same study also demonstrated that L-selectin binding to PECAM-1 is dependent on TNF-α, which is absent in iRhom2-deficient mice. Thus, an additional plausible explanation for the impaired neutrophil infiltration is a dysregulation of cytokine production and neutrophil chemoattractant release, which was therefore also investigated in this study.

Indeed, iRhom2 deficiency resulted in reduced levels of the proinflammatory IL-6 and the neutrophil-attracting chemokine CXCL1 (also known as KC), the murine homolog of IL-8 [32]. However, neither IL-6 nor CXCL1 are among the more than 80 substrates described for ADAM17 [33, 34]. Hence, the release of these mediators may be indirectly regulated by TNF-α, a well-established substrate of ADAM17, whose shedding is enhanced by increased ADAM17 activity following iRhom2 upregulation during inflammation. In fact, TNF-α signaling activates NFκB, thereby promoting the transcription of numerous proinflammatory mediators including TNF-α, IL-6, and IL-8 [35–37]. This pathway is also directly triggered by LPS binding to the Toll-like receptor 4 (TLR4), which explains the observed increase in *Tnf-α* mRNA levels and the release of IL-6 and CXCL1 even in *iRhom2*^-/-^ mice upon LPS stimulation [38]. Released IL-6 in turn can bind to soluble IL-6Rα, generated through ADAM17-mediated shedding, thereby amplifying the inflammatory response via gp130-dependent (trans-)signaling and downstream STAT3 activation [39, 40]. Given that TNF-α is released early during inflammation, it is plausible that ADAM17/iRhom2-dependent TNF-α shedding indirectly regulates the subsequent production of IL-6/IL-6Rα and CXCL1 at later time points.

As the cytokine and chemokine release in the isolated perfused lung (IPL) model where circulating neutrophils are absent still exhibits iRhom2 dependency, resident macrophages are likely critical mediators of the ADAM17-driven inflammatory response. This notion is further supported by the observation that bone marrow derived macrophages (BMDMs) display iRhom2-dependent IL-6 and CXCL1 release following LPS stimulation. Given that macrophages are the primary source of TNF-α, they may act as key regulators during lung inflammation [41]. However, alveolar epithelial cells might also contribute to IL-6 production in response to TNF-α signaling, as previously described [42]. In this study, no differences in the activation of downstream signaling pathways, such as phosphorylated NFκB (pNFκB) or STAT3 (pSTAT3), were detected between LPS-stimulated *iRhom2*^+/+^ and *iRhom2*^-/-^ lung tissue or macrophages. While STAT3 phosphorylation increased upon LPS stimulation, likely due to sufficient IL-6 release in both genotypes, NFκB phosphorylation, which is typically directly induced by LPS signaling, remained unchanged across conditions. This suggests that the selected time points may not have captured transient NFκB activation. Therefore, further studies are required to more precisely define the iRhom2-dependent regulatory mechanisms controlling IL-6 and CXCL1 release.

Taken together, our findings establish iRhom2 as a key regulator of ADAM17 activity in lung inflammation, controlling both neutrophil recruitment and cytokine production. Beyond its direct role in the shedding of proinflammatory cytokines and their receptors, ADAM17 activity may exert broader regulatory effects on inflammatory networks via TNF-α-dependent pathways. Targeting ADAM17 may therefore be a valuable therapeutic strategy in lung injury. Whereas global inhibition of ADAM17 is limited by substantial side effects (reviewed in [43]), selective targeting of iRhom2 may offer a more refined therapeutic approach, dampening pathological inflammation while preserving basal ADAM17 function through the presence of iRhom1.

## Declarations

### Ethics approval and consent to participate

Animal model experiments were approved by local government authorities and were performed according to the German animal protection law (approval-ID: 81-02.04.2019.A271; A4-90029A4). Mice with iRhom2-knockout (*iRhom2^-/-^*) and wildtype (*iRhom2^+/+^*) littermates in a C57BL/6 background derived from in house breeding [20].

## Consent for publication

Not applicable

## Availability of data and materials

The datasets and materials generated during the current study are available from the corresponding author upon reasonable request. Data reanalysis was performed from GSE196638.

## Author Contributions

C.L. S.K., S.D., and A.L. designed the study. C.M. provided methodology. P.K. and R.S. generated the transgenic mouse line. C.L. S.K. and K.S. performed experiments. S.D., S.K., and C.L. analyzed data and performed statistical analysis. A.B. performed bioinformatic analysis. C.L. and A.L. wrote the manuscript. All authors have read and agreed to the entire manuscript.

## Acknowledgments

The authors thank Tanja Woopen, Regina Ben Hamza and Irina Rachimow for perfect technical assistance.

Declaration of generative AI and AI-assisted technologies in the writing process

During the preparation of this work the authors used ChatGPT in order to improve the English language and readability of the manuscript. After using this tool/service, the authors reviewed and edited the content as needed and take full responsibility for the content of the published article.

## Abbreviations

ADAM: a disintegrin and metalloproteinase
ALI: acute lung injury
APC: allophycocyanin
ARDS: acute respiratory distress syndrome
AT1/2: alveolar type 1/2
BAL: bronchoalveolar lavage
BMDM: bone marrow derived macrophages
BSA: bovine serum albumin
CD62L: L-selectin
CXCL1: C-X-C motif chemokine ligand 1
Eef2: eukaryotic elongation factor 2
EL: emphysema lung
ELISA: enzyme-linked immunosorbent assay
FC: flow cytometry
FITC: fluorescein
For: forward
GAPDH: glyceraldehyde 3-phosphate dehydrogenase
HL: healthy lung
IL-(6): interleukin-(6)
IL-6Rα: interleukin-6 receptor α
IPL: isolated perfused lungs
LPS: lipopolysaccharide
(p)NFκB: (phosphorylated) nuclear factor kappa B
PBS: phosphate-buffered saline
PE: phycoerythrin
PECAM-1: platelet endothelial cell adhesion molecule-1
PFA: paraformaldehyde
Pla2: phospholipase A2
qPCR: quantitative real-time polymerase chain reaction
Rev: reverse
RHBDF/iRhom: inactive rhomboid protein family
Rpl13: ribosomal protein L13
Rps29: ribosomal protein S29
SDS: sodium dodecyl sulfate
(p)STAT3: (phosphorylated) signal transducer and activator of transcription 3
SMC: smooth muscle cell
Tbp: TATA-box binding protein
TEM: transendothelial migration
TLR4: toll-like receptor 4
TNF-α: tumor necrosis factor-α
W/D ratio: wet-to-dry ratio

---

**Suppl. Figure S 1:**
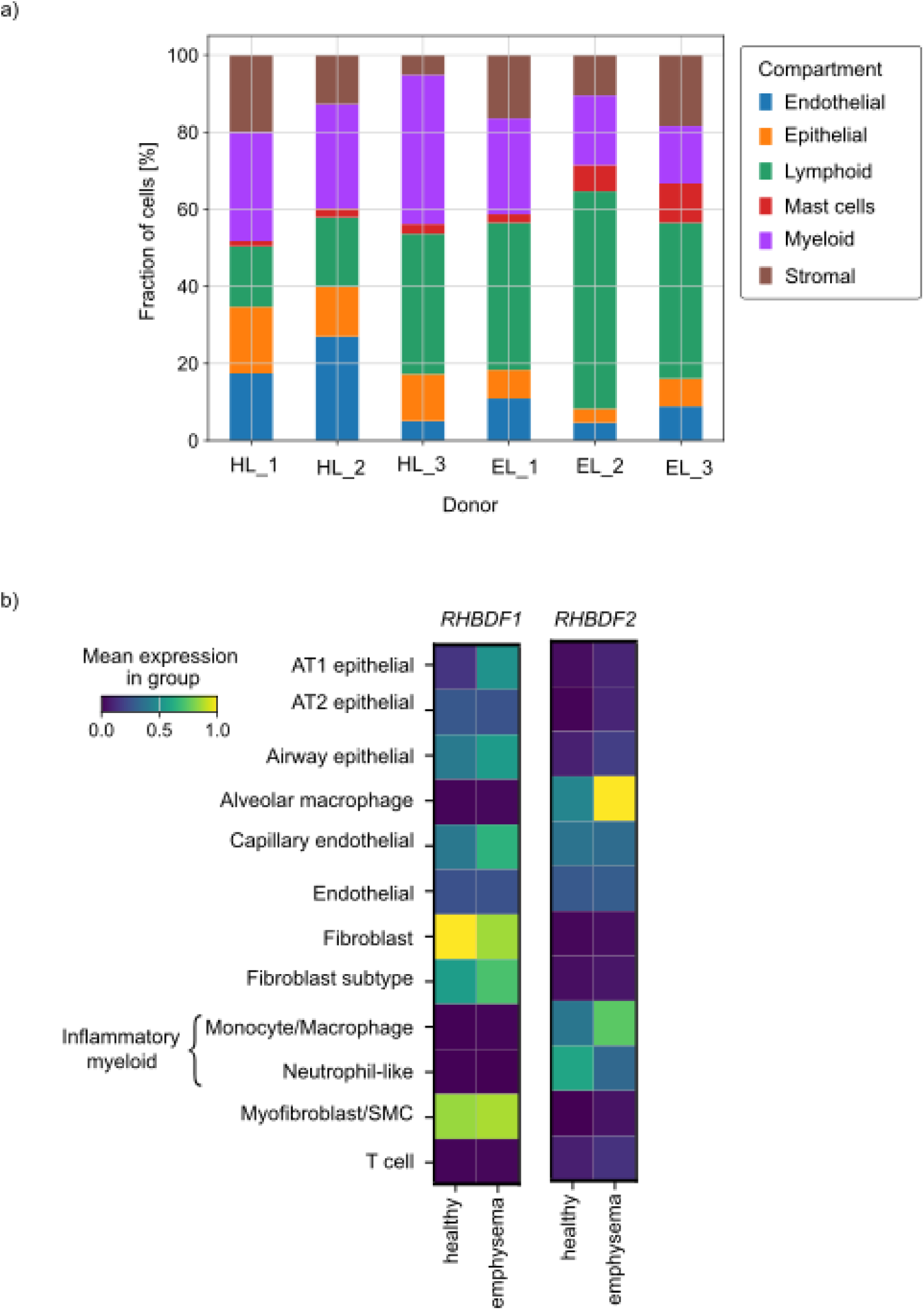
Single cell mRNA sequencing of emphysema patients. **a)** Distribution of cell types in biopsies of healthy and emphysema lungs. Data derived from GSE196638. HL – healthy lung; EL – emphysema lung. **b)** Heat map showing mRNA expression of *RHBDF1* and *RHBDF2* in different cell types of healthy and emphysema lung biopsies.

**Suppl. Figure S 2:**
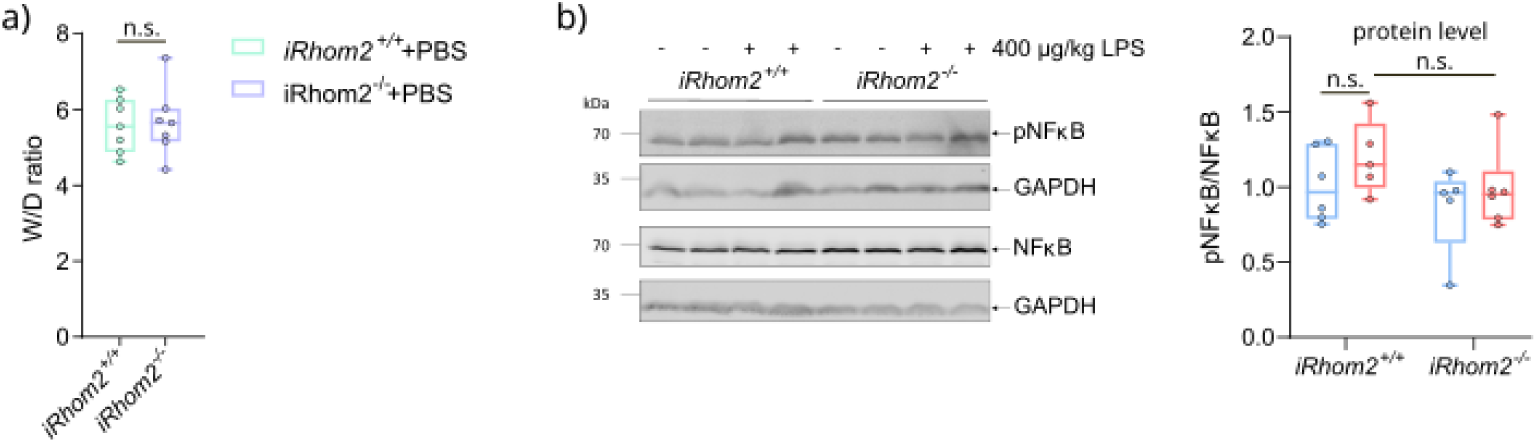
iRhom2 deficiency does not influence wet-to-dry ratio in control experiments and NFκB phosphorylation in the isolated perfused lung model. **a)** Wet-to-dry (W/D) ratio of IPL-derived lungs of *iRhom2^+/+^* and *iRhom2^-/-^* mice subjected to 50 µl PBS. The ratio was calculated by dividing the lung mass dried for 72 h at 60 °C by their wet mass immediately after the experiment. **b)** Western blot analysis of phosphorylated NFκB (pNFκB) and NFκB from processed lung tissue. GAPDH serves as a loading control. Quantified data show the determined protein ratio of pNFκB and the corresponding NFκB. **a-b)** Quantitative data are shown as Tukey box plots from at least five independent experiments. Statistical analysis was performed using a generalized linear mixed model with a post-hoc false discovery rate (FDR) correction for multiple comparisons. n.s. – not significant.

**Suppl. Figure S 3:**
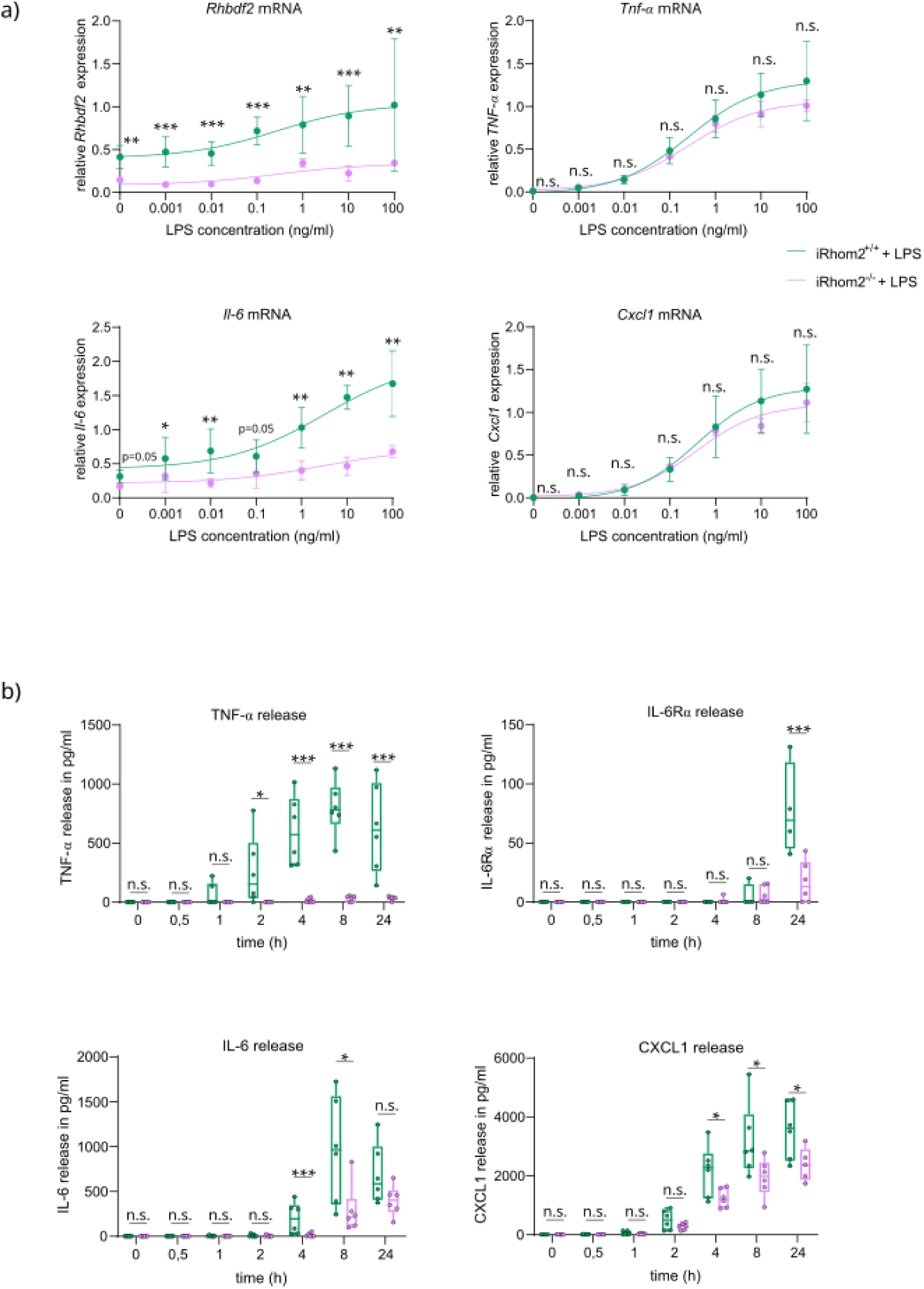
Different LPS concentrations and time points lead to iRhom2-dependent inflammatory response. **a)** Relative mRNA expression of *Rhbdf2*, *Tnf-α, Cxcl1* and *Il-6* in *iRhom2^+/+^* and *iRhom2^-/-^* BMDMs after stimulation with 0, 0.001, 0.01, 0.1, 1, 10 and 100 ng/ml LPS for 1 h. Data are represented as mean ± SD from three independent experiments. **b)** Protein release of TNF-α, CXCL1, IL-6 and IL-6Rα in pg/ml measured by ELISA with supernatant from *iRhom2^+/+^* and *iRhom2^-/-^* BMDMs after 1 ng/ml LPS stimulation for 0, 0.5, 1, 2, 4, 8 and 24 h. Data are shown as Tukey box plots from at least four independent experiments. **a-b)** Statistical analysis was performed using a generalized linear mixed model with a post-hoc false discovery rate (FDR) correction for multiple comparisons. Statistical differences are indicated by bars and asterisks with * p ≤ 0.05, ** p ≤ 0.01 and *** p ≤ 0.001, n.s. – not significant.

## References

1. Eisner, M.D., et al., Efficacy of Low Tidal Volume Ventilation in Patients with Different Clinical Risk Factors for Acute Lung Injury and the Acute Respiratory Distress Syndrome. American Journal of Respiratory and Critical Care Medicine, 2001. 164(2): p. 231–236.

2. Matthay, M.A. and G.A. Zimmerman, Acute lung injury and the acute respiratory distress syndrome: four decades of inquiry into pathogenesis and rational management. Am J Respir Cell Mol Biol, 2005. 33(4): p. 319–27.

3. Thompson, B.T., C. Chambers Rachel, and D. Liu Kathleen, Acute Respiratory Distress Syndrome. New England Journal of Medicine. 377(6): p. 562–572.

4. Abraham, E., et al., Neutrophils as early immunologic effectors in hemorrhage-or endotoxemia-induced acute lung injury. American Journal of Physiology-Lung Cellular and Molecular Physiology, 2000. 279(6): p. L1137–L1145.

5. Meduri, G.U., et al., Inflammatory Cytokines in the BAL of Patients With ARDS: Persistent Elevation Over Time Predicts Poor Outcome. CHEST, 1995. 108(5): p. 1303–1314.

6. Li, T., et al., Tumor Necrosis Factor-α Plays an Initiating Role in Extracorporeal Circulation-induced Acute Lung Injury. Lung, 2013. 191: p. 207–214.

7. Black, R.A., et al., A metalloproteinase disintegrin that releases tumour-necrosis factor-alpha from cells. Nature, 1997. 385(6618): p. 729–733.

8. Mauer, J., et al., Signaling by IL-6 promotes alternative activation of macrophages to limit endotoxemia and obesity-associated resistance to insulin. Nat Immunol, 2014. 15(5): p. 423–30.

9. Horiuchi, K., et al., Cutting Edge: TNF-α-Converting Enzyme (TACE/ADAM17) Inactivation in Mouse Myeloid Cells Prevents Lethality from Endotoxin Shock1. The Journal of Immunology, 2007. 179(5): p. 2686–2689.

10. Kuebler, W.M., et al., Role of l-Selectin in Leukocyte Sequestration in Lung Capillaries in a Rabbit Model of Endotoxemia. American Journal of Respiratory and Critical Care Medicine, 2000. 161(1): p. 36–43.

11. Aljohmani, A., et al., Leukocytic ADAM10 and ADAM17 modulate disease severity and systemic outcome in bacterial and viral pneumonia. Eur Respir J, 2026. 67(2).

12. Dreymueller, D., S. Uhlig, and A. Ludwig, ADAM-family metalloproteinases in lung inflammation: potential therapeutic targets. American Journal of Physiology-Lung Cellular and Molecular Physiology, 2015. 308(4): p. L325–L343.

13. Giese, A.A., et al., Inflammatory activation of surface molecule shedding by upregulation of the pseudoprotease iRhom2 in colon epithelial cells. Scientific Reports, 2021. 11(1): p. 24230.

14. Issuree, P.D.A., et al., iRHOM2 is a critical pathogenic mediator of inflammatory arthritis. The Journal of Clinical Investigation, 2013. 123(2): p. 928–932.

15. Christova, Y., et al., Mammalian iRhoms have distinct physiological functions including an essential role in TACE regulation. The EMBO Reports, 2013. 14(10): p. 884–890.

16. Adrain, C., et al., Tumor Necrosis Factor Signaling Requires iRhom2 to Promote Trafficking and Activation of TACE. Science, 2012. 335(6065): p. 225–228.

17. Veit, M., et al., Mutagenesis of the ADAM17-phosphatidylserine-binding motif leads to embryonic lethality in mice. Life Sci Alliance, 2019. 2(5).

18. McIlwain, D.R., et al., iRhom2 regulation of TACE controls TNF-mediated protection against Listeria and responses to LPS. Science, 2012. 335(6065): p. 229–32.

19. Wang, C., et al., Dysregulated lung stroma drives emphysema exacerbation by potentiating resident lymphocytes to suppress an epithelial stem cell reservoir. Immunity, 2023. 56(3): p. 576–591.e10.

20. Düsterhöft, S., et al., The iRhom homology domain is indispensable for ADAM17-mediated TNFα and EGF receptor ligand release. Cell Mol Life Sci, 2021. 78(11): p. 5015–5040.

21. Dreymueller, D., et al., Lung endothelial ADAM17 regulates the acute inflammatory response to lipopolysaccharide. EMBO Molecular Medicine, 2012. 4(5): p. 412–423.

22. Held, H.D., C. Martin, and S. Uhlig, Characterization of airway and vascular responses in murine lungs. British Journal of Pharmacology, 1999. 126.

23. Krabbe, J., et al., The effects of hydroxyethyl starch and gelatine on pulmonary cytokine production and oedema formation. Scientific Reports, 2018. 8(1): p. 5123.

24. Pruessmeyer, J., et al., Leukocytes require ADAM10 but not ADAM17 for their migration and inflammatory recruitment into the alveolar space. Blood, 2014. 123(26): p. 4077–4088.

25. Degasperi, A., et al., Evaluating Strategies to Normalise Biological Replicates of Western Blot Data. PLOS ONE, 2014. 9(1): p. e87293.

26. Bell, J.H., et al., Role of ADAM17 in the ectodomain shedding of TNF-α and its receptors by neutrophils and macrophages. Journal of Leukocyte Biology, 2007. 82(1): p. 173–176.

27. Lorenzen, I., et al. Control of ADAM17 activity by regulation of its cellular localisation. Scientific reports, 2016. 6, 35067 DOI: 10.1038/srep35067.

28. Skurski, J., et al., The Threshold Effect: Lipopolysaccharide-Induced Inflammatory Responses in Primary Macrophages Are Differentially Regulated in an iRhom2-Dependent Manner. Front Cell Infect Microbiol, 2020. 10: p. 620392.

29. Yang, S.-C., et al., Understanding the role of neutrophils in acute respiratory distress syndrome. Biomedical Journal, 2021. 44(4): p. 439–446.

30. Arndt, P.G., et al., Leukocyte ADAM17 regulates acute pulmonary inflammation. PLoS One, 2011. 6(5): p. e19938.

31. Rahman, I., et al., L-selectin regulates human neutrophil transendothelial migration. Journal of Cell Science, 2021. 134(3).

32. Sawant, K.V., et al., Chemokine CXCL1 mediated neutrophil recruitment: Role of glycosaminoglycan interactions. Scientific Reports, 2016. 6(1): p. 33123.

33. Scheller, J., et al., ADAM17: a molecular switch to control inflammation and tissue regeneration. Trends in Immunology, 2011. 32(8): p. 380–387.

34. Zunke, F. and S. Rose-John, The shedding protease ADAM17: Physiology and pathophysiology. Biochimica et Biophysica Acta (BBA) - Molecular Cell Research, 2017. 1864(11, Part B): p. 2059–2070.

35. Aggarwal, B.B., Signalling pathways of the TNF superfamily: a double-edged sword. Nature Reviews Immunology, 2003. 3(9): p. 745–756.

36. Wang, Y.H., et al., TNFα induced IL-8 production through p38 MAPK-NF-kB pathway in human hepatocellular carcinoma cells. Zhonghua Gan Zang Bing Za Zhi, 2011. 19(12): p. 912–6.

37. Kurokouchi, K., et al., TNF-alpha increases expression of IL-6 and ICAM-1 genes through activation of NF-kappaB in osteoblast-like ROS17/2.8 cells. J Bone Miner Res, 1998. 13(8): p. 1290–9.

38. Luo, R., et al., An examination of the LPS-TLR4 immune response through the analysis of molecular structures and protein-protein interactions. Cell Commun Signal, 2025. 23(1): p. 142.

39. Hibi, M., et al., Molecular cloning and expression of an IL-6 signal transducer, gp130. Cell, 1990. 63(6): p. 1149–1157.

40. Aliyu, M., et al., Interleukin-6 cytokine: An overview of the immune regulation, immune dysregulation, and therapeutic approach. International Immunopharmacology, 2022. 111: p. 109130.

41. Xiao, J., et al., Monocyte-Derived Macrophages Induce Alveolar Macrophages Death via TNF-α in Acute Lung Injury. Immun Inflamm Dis, 2024. 12(12): p. e70081.

42. Yan, C., et al., TNF-α induction of IL-6 in alveolar type II epithelial cells: Contributions of JNK/c-Jun/AP-1 element, C/EBPδ/C/EBP binding site and IKK/NF-κB p65/κB site. Molecular Immunology, 2018. 101: p. 585–596.

43. Calligaris, M., et al., Strategies to Target ADAM17 in Disease: From its Discovery to the iRhom Revolution. Molecules, 2021. 26(4).

